# The photoactive site modulates current rectification and channel closing in the natural anion channelrhodopsin *Gt*ACR1

**DOI:** 10.1101/630327

**Authors:** Oleg A. Sineshchekov, Elena G. Govorunova, Hai Li, Xin Wang, John L. Spudich

**Affiliations:** Center for Membrane Biology, Department of Biochemistry and Molecular Biology, University of Texas Health Science Center – McGovern Medical School, Houston, TX 77030, USA

## Abstract

The crystal structure of *Gt*ACR1 from *Guillardia theta* revealed an intramolecular tunnel predicted to expand to form the anion-conducting channel upon photoactivation (Li et al. 2019). The location of the retinylidene photoactive site within the tunnel raised the question of whether, in addition to triggering channel opening by photoisomerization, the site also participates in later channel processes. Here we demonstrate the involvement of the photoactive site in chloride conductance and channel closing. Electrostatic perturbation of the photoactive retinylidene Schiff base region by glutamate substitutions alters the rectification of the photocurrent as well as channel closing kinetics. Substitutions on opposite sides of the photoactive site causes opposite changes, with channel closing kinetically correlated with Schiff base deprotonation, and the extent of these effects closely correlate with distance of the introduced glutamyl residue from the photoactive site.

## Introduction

Anion channelrhodopsins from the cryptophyte alga *Guillardia theta* (*Gt*ACRs) are the most potent optogenetic inhibitors of neuronal firing and regulators of animal behavior currently available (Govorunova et al. 2015; Mahn et al. 2018; Messier et al. 2018; Mohammad et al. 2017; Wilson et al. 2018). Recently we and others have obtained high-resolution X-ray structures of the dark (closed) state of *G. theta* anion channelrhodopsin 1 (*Gt*ACR1) (Kim et al. 2018; Li et al. 2019). In contrast to available structures of cation channelrhodopsins (CCRs) from green algae (Kato et al. 2012; Oda et al. 2018; Volkov et al. 2017), *Gt*ACR1 exhibits a narrow continuous intramolecular tunnel formed by helices 1-3 and 7 that connects the cytoplasmic and extracellular aqueous phases and presumably expands upon illumination to pass anions (Li et al. 2019). However, no open channel structure is yet available, and many questions regarding its architecture remain unanswered. In particular, a possible role of the second extracellular vestibule (Kim et al. 2018) in anion conductance has not so far been tested experimentally. Also, the functional significance of the many structural differences between ACRs and CCRs is not yet clear.

The voltage dependence of the peak photocurrent generated by *Gt*ACR1 is virtually linear when tested with symmetrical concentrations of the permeant ion species on the two sides of the membrane (Govorunova et al. 2015). However, kinetic analysis of single-turnover laser flash-evoked photocurrents revealed two components with current-voltage relationships that show rectification (i.e., deviation from linearity) in the opposite directions (Sineshchekov et al. 2015). The amplitude of the fast component of the photocurrent decay exhibited outward rectification, whereas that of the slow component showed inward rectification. Both components of channel closing correlate with deprotonation of the Schiff base in the proposed conductive intermediate L (Sineshchekov et al. 2016). Depletion of L proceeds in two kinetics phases: a fast reversible L⇔M transition and the slower irreversible depletion of M and hence depletion of L. These phases are readily measured spectroscopically by monitoring the accumulation and dissipation of the far blue-shifted M intermediate.

We conducted a protein-wide carboxyl residue substitution screen to study the functional consequences of mutagenetic modifications of the protein’s electrostatic profile with pulsed continuous light stimulation. As reported here, our screen revealed that mutations only of the residues contributing to the intramolecular tunnel that we had detected in the closed-state crystal structure of *Gt*ACR1 (Li et al. 2019) caused strong changes in rectification, which confirmed our hypothesis that the pathway for anion passage is indeed created by expansion of this tunnel upon photoexcitation. The acidic substitutions of the tunnel residues located to the extracellular side of the retinylidene photoactive site cause inward rectification, whereas those of the residues located to the cytoplasmic side of the photoactive site confer outward rectification.

To gain further insight into the mechanism of channel closing and rectification in *Gt*ACR1 we performed quantitative kinetic analysis of laser flash-evoked photocurrents in several mutants with glutamates introduced on the extracellular and cytoplasmic sides of the photoactive site. The data showed opposite effects on both the relative amplitudes and the rates of the two channel closing components in these two groups of mutants. This observation and the dependence of the effects on the distance between the photoactive site and the introduced negative charge strongly indicate that oppositely directed charge movements involving the photoactive site are involved in fast and slow channel closing. The opposite charge movements would explain the opposite rectification in the wild type *Gt*ACR1. Moreover, we observed that in the mutants as in the wild type, both fast and slow closing of the channel correlate with deprotonation of the Schiff base in the proposed conductive intermediate L.

## Results and Discussion

### Photocurrents from the Ala61 and Ala75 mutants under continuous light stimulation

Two of the four Glu residues conserved in the second transmembrane helix (TM2) of CCRs are neutral residues in ACRs, most frequently Ala (Figure 1A and B). In *Gt*ACR1 these are Ala61 in the cytoplasmic end of the intramolecular tunnel detected in the dark structure of *Gt*ACR1, and Ala75 in the extracellular end. Previously we had shown that substitution of Glu for either Ala61 or Ala75 led to strong inhibition of photocurrents in *Gt*ACR1 (Li et al. 2019). These mutations also caused strong rectification of the voltage dependence of the peak photocurrent (i.e., the current measured at the positive voltage differed from that of the negative voltage of the same absolute value). The direction of rectification was outward in the A61E mutant and inward in the A75E mutant (Figure 1C and D). Substitution of Asp for Ala61 or Ala75 caused similar effects as substitution of Glu at the corresponding positions (Figure 1C), whereas mutations of Ala61 to Ser or Thr produced no effect on the shape of the current-voltage relationship (Figure 1 – figure supplement 1). These results indicate that the changes in rectification observed in the mutants are brought about by the negative charge of the mutated side chain rather than by its length.

**Figure 1.**
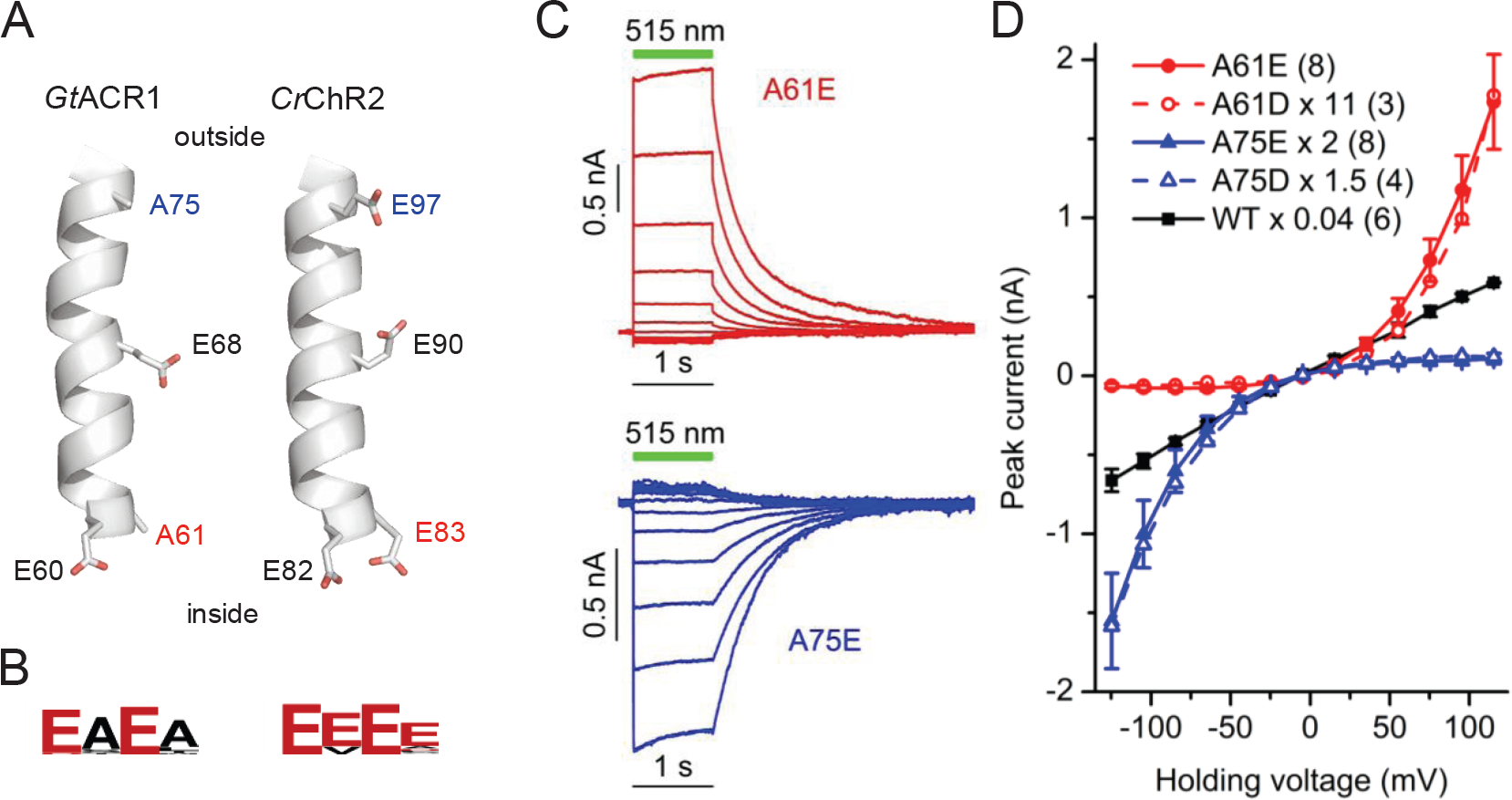
Asp/Glu substitutions at positions 61 and 75 change voltage dependence in the opposite directions. (**A**) Structures of TM2 in *Gt*ACR1 (PDB code 6EDQ) and *Chlamydomonas reinhardtii* channelrhodopsin 2 (*Cr*ChR2, PDB code 6EID). The side chains of the Glu residues conserved in CCRs and their counterparts in ACRs are shown as sticks. (**B**) Residue conservation logos of the corresponding positions in all currently known functional ACRs and CCRs. (**C**) Series of photocurrents recorded from *Gt*ACR1_A61E and *Gt*ACR1_A75E mutants expressed in HEK293 cells at the voltages at the amplifier output changed from -120 to 120 mV (bottom to top photocurrent transient in each set) in 20-mV increments. The green bars show duration of illumination with 515 nm light (7.7 mW mm^-2^). For solution compositions and other details see Methods. (**D**) The current-voltage relationships for the wild-type *Gt*ACR1 and its mutants scaled as indicated in the legend. The data points are the mean values ± SE (n = 3-8 cells; n for each variant is indicated in the legend in parentheses).

### Carboxyl residue substitution screen

Measurement of rectification of the current-voltage dependence provides a more specific functional test than measurement of the current amplitude at any single holding voltage, because suppression of the amplitude could be brought about by detrimental effects of the mutations on protein folding, which is difficult to distinguish from direct local effects on channel conductance. We made Glu or Asp substitution mutants of 75 residues in *Gt*ACR1 and determined the rectification index (RI) defined as the ratio of the peak current at 60 mV to that at -60 mV. The mean RI values are shown in Figure 2, and the mean current amplitudes recorded at 60 and -60 mV are shown in Figure 2 – figure supplements 1 and 2, respectively. Using the one-sample Wilcoxon signed rank test at p < 0.01, we classified the mutants into five categories according to their median RI values: RI < 0.5, 0.5 < RI <1, RI = 1, 1 < RI < 2 and RI > 2 (see Figure 2 source data file for numerical data and full statistical analysis). All mutations that caused strong rectification in either direction also reduced the current amplitude, as compared to the wild type, even at the favorable voltage (Figure 2 – figure supplement 3). Glu replacement of Leu64 and Thr101 which, together with Met101, form the central constriction of the tunnel (Li et al. 2019), caused such dramatic reduction of photocurrents that the RI could not be determined.

**Figure 2.**
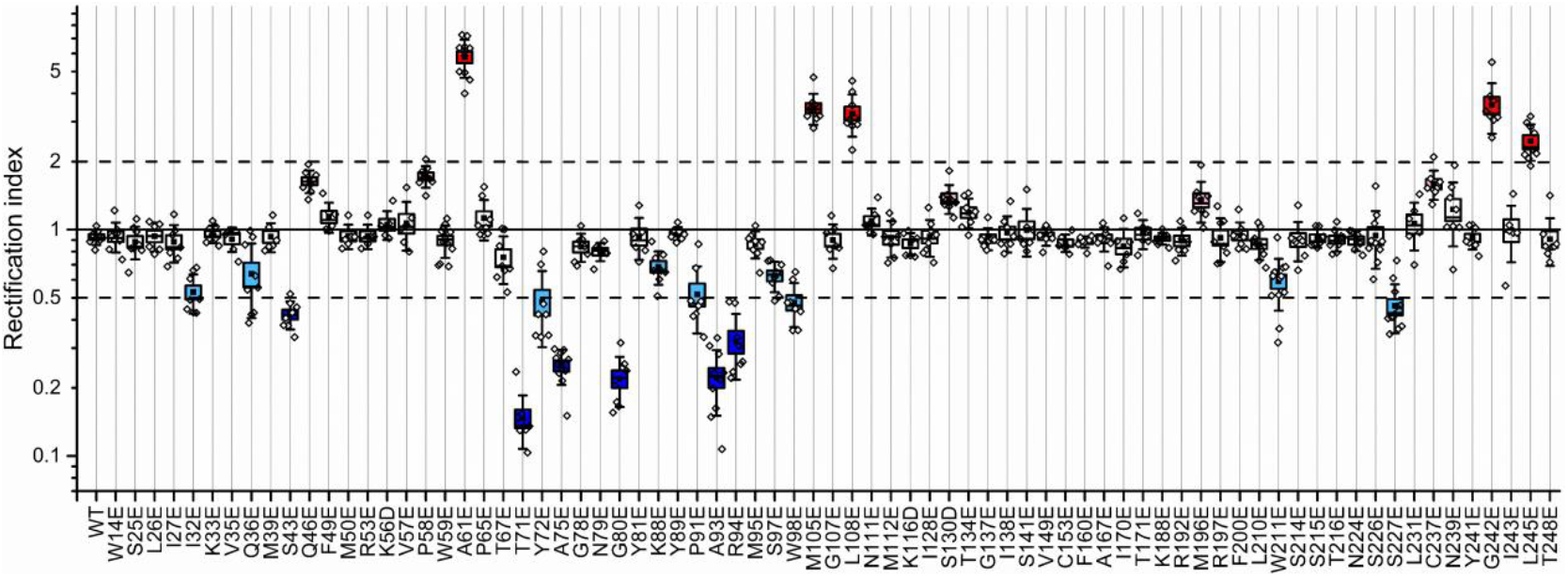
Rectification index in wild-type *Gt*ACR1 and its Glu or Asp substitution mutants. The rectification index was calculated as the ratio of the peak photocurrent recorded at 60 mV to that recorded at -60 mV in response to 1-s pulses of 515 nm light with 131 and 156 mM Cl^-^in the pipette and bath, respectively (for other solution components see Methods). The black squares, mean; line, median; box, SE; whiskers, SD; empty diamonds, raw data recorded from individual cells.

Distribution of the residues mutation of which showed statistically significant rectification within the *Gt*ACR1 molecule displayed a clear pattern (Figure 3A). Strong rectification in either direction (RI < 0.5 or > 2) was observed almost exclusively in the mutants of the residues that line the intramolecular tunnel. These results strongly support our hypothesis that the anion conduction pathway is created in illuminated *Gt*ACR1 by expansion of the intramolecular tunnel we detected in its dark (closed) state structure.

**Figure 3.**
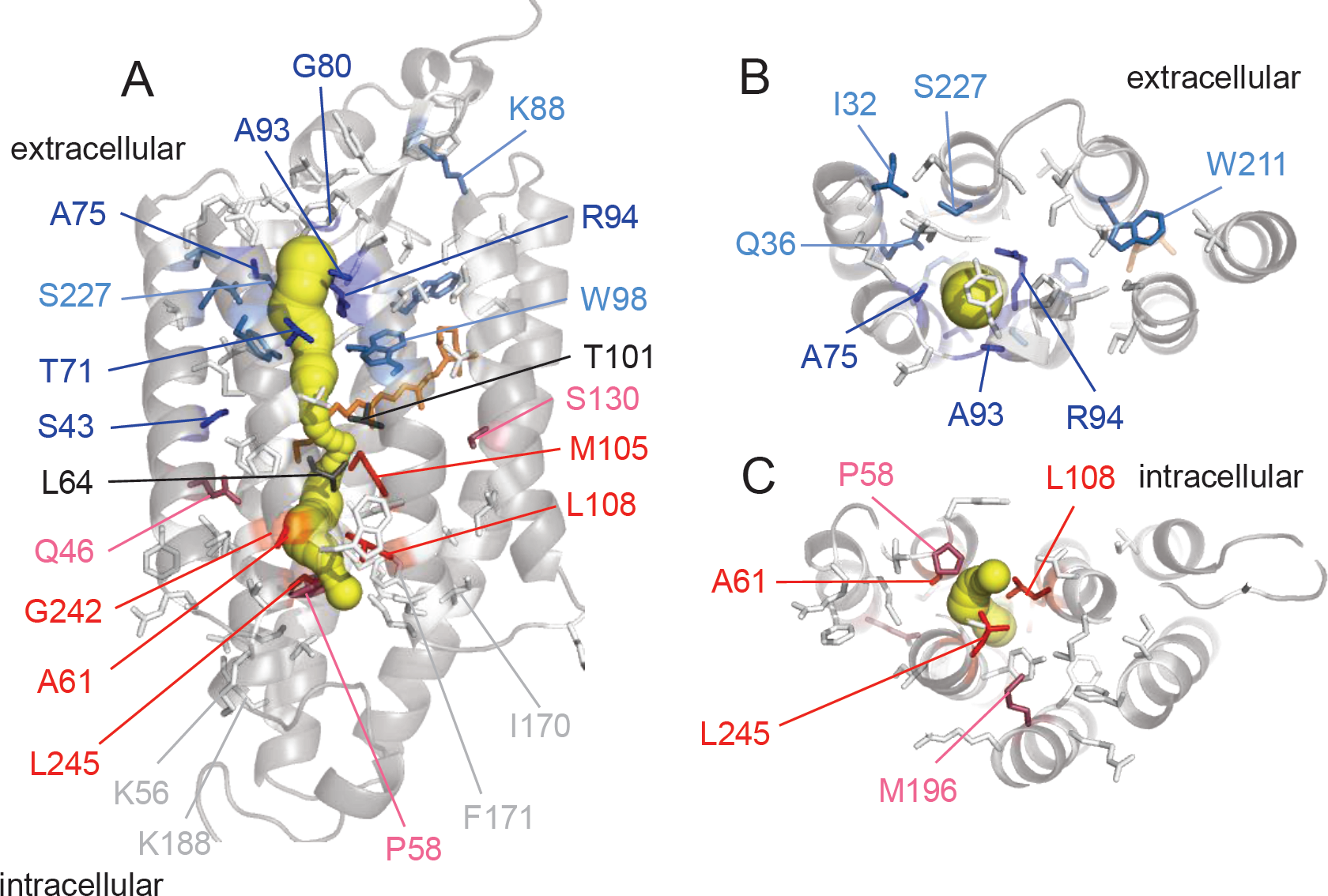
A side (A), top/extracellular (B) and bottom/intracellular (C) views of *Gt*ACR1 structure (6EDQ) showing the side chains of the tested residues color-coded according to the rectification index measured in their mutants. Only 15-Å slabs centered on Ala75 and Ala61, respectively, are shown in (B) and (C) for clarity. The intramolecular tunnel is shown in yellow. The color code for amino acid side chains is as follows: dark blue, RI < 0.5; light blue, 0.5 < RI < 1; light grey, RI = 1; pink, 1 < RI <2; red, RI > 2, as determined by the one-sample Wilcoxon signed rank test at p < 0.01; black, RI could not be determined because of severe suppression of photocurrents.

The direction of rectification observed in the mutants strongly correlated with the position of the mutated residue within the tunnel. The residues, replacement of which with acidic residues caused strong inward rectification (Thr71, Ala75, Gly80, Ala93 and Arg94), form the extracellular portion of the tunnel (Figure 3B); those, replacement of which caused strong outward rectification (Ala61, Met105, Leu108, Gly242 and Leu245) form the cytoplasmic portion (Figure 3C). Of all tested residues outside the tunnel only replacement of Ser43 caused strong rectification (RI < 0.5). Modest rectification (i.e. with RI between 0.5 and 2, but significantly different from 1) resulted from Glu or Asp substitutions of several other residues (including Gln46 and Ser130) that do not contribute to the tunnel in the dark state. Our interpretation is that mutations of these residues either indirectly influence the chloride pathway via long-range interactions or that these residue positions become exposed to the tunnel after its expansion under illumination.

### Probing the role of the second extracellular vestibule

An intramolecular cavity opened to the extracellular space (a vestibule) was found in the *Gt*ACR1 dark structure in addition to the continuous tunnel (Figure 3 – figure supplement 1). The crystal structures of the hybrid CCR known as C1C2 (Kato et al. 2012) and its Cl^-^-conducting mutant iC++ (Kato et al. 2018) also show a vestibule in the corresponding position (EV2), which is thought to represent the main extracellular entry into the channel pore in these molecules. In the dark *Gt*ACR1 structure EV2 is separated from the intramolecular tunnel by hydrogen-bonded Tyr81, Arg94 and Glu223 (Kim et al. 2018; Li et al. 2019). However, it could not be excluded that upon photoexcitation this hydrogen-bonded network is broken and the second vestibule is merged with the tunnel. To test this hypothesis, we measured the RI in the mutants of the residues that line the second vestibule. Individual glutamate replacement of nearly all of these residues (Pro91, Met95, Ser214, Ser 215, Thr216 and Ser226; Figure 3 – figure supplement 1) did not affect rectification, which suggests that the second extracellular vestibule plays only a minor, if any, role in anion conduction in *Gt*ACR1. Thus our functional data provide empirical confirmation of the stability of hydrogen bonding between Tyr81, Arg94 and Glu223 deduced from all-atom molecular dynamic simulations of *Gt*ACR1 (Kato et al. 2018).

### Photocurrents and absorption changes under single-turnover excitation

Illumination with pulses of continuous light is not suitable for correlation of the kinetics of channel opening and closing with the laser-flash induced photocycle data. The main limitation is that a pulse of continuous light produces a mixture of photocycle intermediates, the composition of which depends on the intensity, duration and wavelength of light. In wild type *Gt*ACR1 rectification was observed in laser flash-induced currents, but under pulse illumination the rectification is eliminated by the superposition of two kinetics components with opposite rectification (Sineshchekov et al. 2015). Therefore, analysis of the channel mechanisms requires measurements of photocurrents under single-turnover conditions, i.e. laser flash excitation. To evaluate the possible influence of rectifying mutations on the individual current components, we measured laser flash-induced photocurrents from six rectifying mutants selected for an optimal combination of their RI values and photocurrent amplitude (Figure 2 – figure supplement 3). The initial analyzed set comprised three outwardly rectifying mutants (A61E, G242E and Q46E) and three inwardly rectifying mutants (A75E, G80E and S227E).

Substitution with a glutamyl residue caused a strong perturbation of the kinetics of single-turnover laser flash-induced currents in all six analyzed mutants (Figure 4). In each mutant channel closing proceeded in two kinetically different phases, as in the wild type. We performed multiexponential fitting of the current traces and obtained the times constants (τ) and amplitudes of the two components of the photocurrent decay. At -60 mV, the ratio of the τ values of slow to fast closing increased from 20 to 1,000-fold in mutants, compared to the wild type (Figure 4 – figure supplement 1). In all mutants, the amplitude of the fast component showed no or outward rectification (Figure 4 – figure supplement 2), as in the wild type (Sineshchekov et al. 2015). However, the slow component showed strong inward rectification only in inwardly rectifying

**Figure 4.**
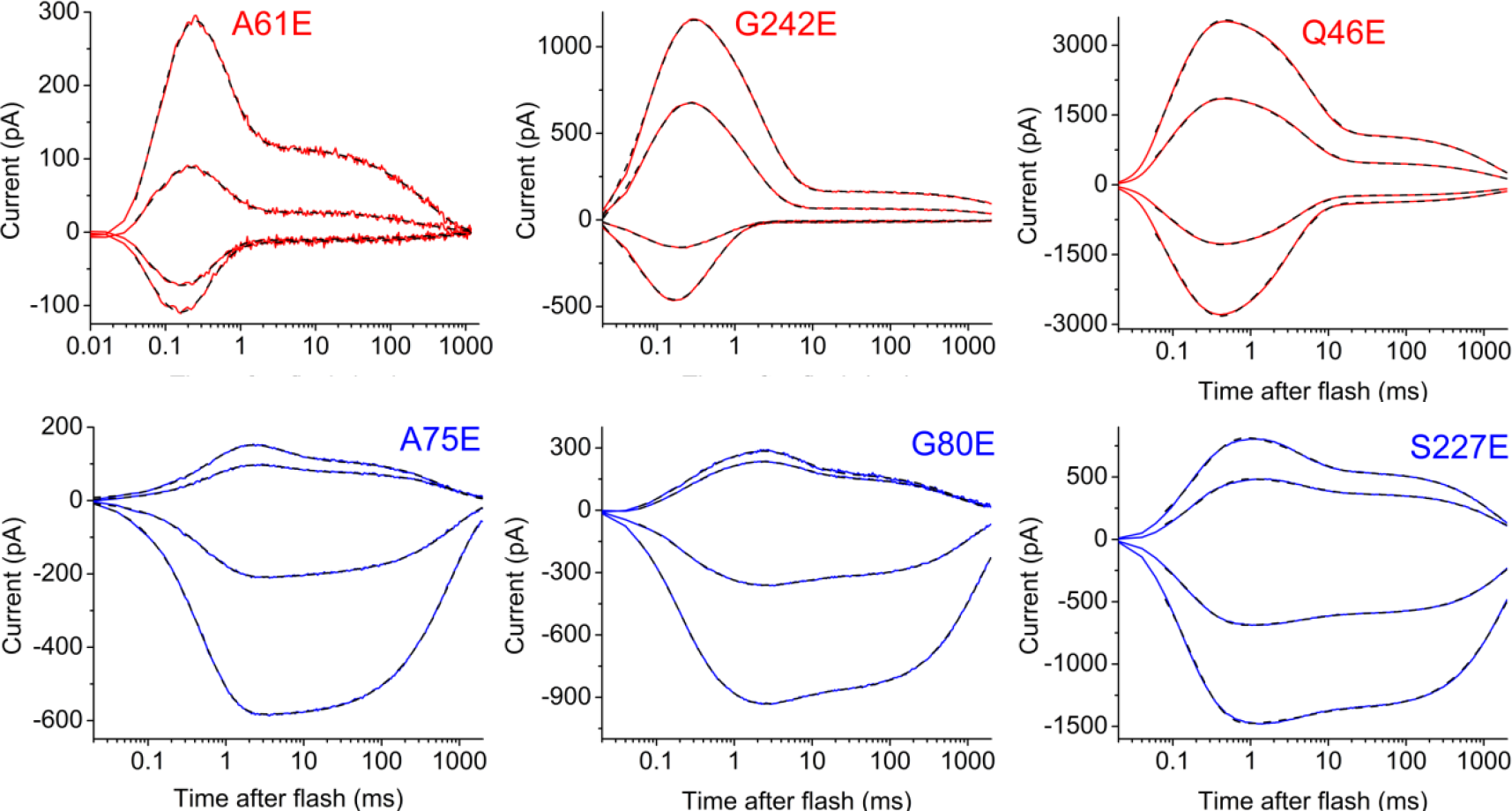
Photocurrent traces recorded from the rectifying *Gt*ACR1 mutants upon excitation with laser flashes. The current traces were recorded at -60, -30, 30 and 60 mV at the amplifier output (from bottom to top). The solid colored lines show the experimental data, the dashed black lines, their multiexponential fits.

Next, we analyzed contributions of the amplitude of the fast component to the overall current (its ratio to the sum of the two components). At -60 mV this contribution was roughly an order of magnitude greater in outwardly rectifying mutants than in inwardly rectifying mutants, while in the wild type the amplitudes of the components were almost equal (Figure 5A). The rate of the fast closing increased in all mutants, as compared to the wild type, but the rate of fast closing in outwardly rectifying mutants was also almost an order of magnitude greater than that in inwardly rectifying mutants (Figure 5B). Upon shifting of the holding voltage to more positive values, the contribution of the fast phase decreased in the outwardly rectifying mutants and increased in the inwardly rectifying mutants (Figure 5C). In summary, introduction of glutamate inwardly to the photoactive site activated fast closing of the channel which has outward rectification, while placement of the negative charge outwardly to the photoactive site shifted the closing in favor of the slow component with inward rectification (Figure 5, Figure 4 – figure supplement 2).

**Figure 5.**
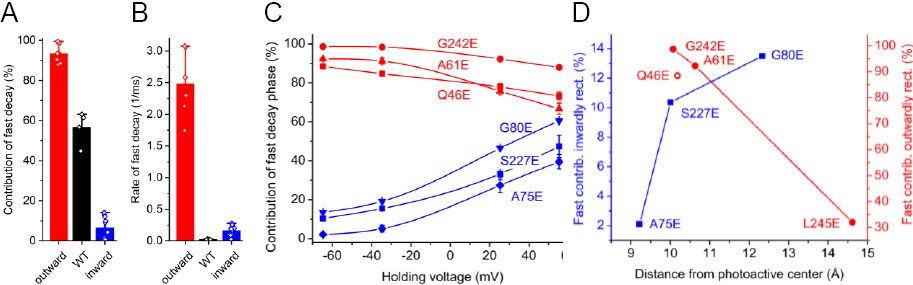
Influence of the rectifying mutations on the kinetics of laser-flash induced photocurrents. (**A** and **B**) The mean contributions (A) and time constants (B) of the fast current decay phase in the outwardly rectifying mutants, wild type and inwardly rectifying mutants. (**C**) The voltage dependence of the contribution of the fast decay phase in the individual rectifying mutants. The data points are the means values ± SE (n = 3-4 cells). (**D**) The dependence of the contribution of the fast decay phase on the distance from the side chain of the mutated residue to the photoactive site.

We hypothesized that the opposite effects of an acidic group placed outward or inward of the photoactive site on the channel kinetics was due to its electrostatic influence on the site (comprising the protonated Schiff base and two potential proton acceptors, Glu68 and Asp234 (Sineshchekov et al. 2015; Sineshchekov et al. 2016)). To test this hypothesis, we made homology models of the mutants (Figure 5 – figure supplement 1) and calculated the mean distances from the introduced Glu side chain to the Schiff base, Glu68 and Asp234 for each mutant. The mean distances were very similar in the A61E, G242E and Q46E mutants, so we added to our analysis a fourth outwardly rectifying mutant, L245E, in which the introduced glutamate is located farther away from the photoactive site. A series of laser flash-evoked current traces generated by *Gt*ACR1_L245E is shown in Figure 5 – figure supplement 2A. In this mutant, the contribution of the fast decay phase at -60 mV was smaller than in the other three outwardly rectifying mutants analyzed, consistent with its more distant location, but its voltage dependence was similar to that in the other outwardly rectifying mutants (Figure 5 – figure supplement 2B). In each group of the mutants, outwardly and inwardly rectifying, the contribution of the fast phase was proportional to the distance of the mutated residue from the photoactive site (Figure 5D). The contribution measured in the Q46E mutant was smaller than expected from the monotonic proportionality, which may be explained by considering that this residue is located outside of the tunnel, unlike the other residues analyzed, but may be inside the channel in the open state (Figure 3A).

To probe the influence of the introduced acidic group on the photoactive site, we expressed the A61E and A75E mutants in *Pichia* and carried out measurements of laser flash-induced absorbance changes. In both these mutants we observed temporal correlation between Schiff base deprotonation (M formation) and the fast channel closing, and between Schiff base reprotonation (M decay) and slow channel closing (Figure 6A and C), as we have reported earlier in the wild-type *Gt*ACR1 (Sineshchekov et al. 2016). The outwardly rectifying A61E mutation accelerated both M formation and fast channel closing, whereas the inwardly rectifying A75E mutation slowed both of these processes. Absorption change differences between the two mutants were also observed at other characteristic wavelengths (Figure 6B and D). These results confirm that the influence of the rectifying mutations on channel kinetics results from perturbation of the photoactive site.

**Figure 6.**
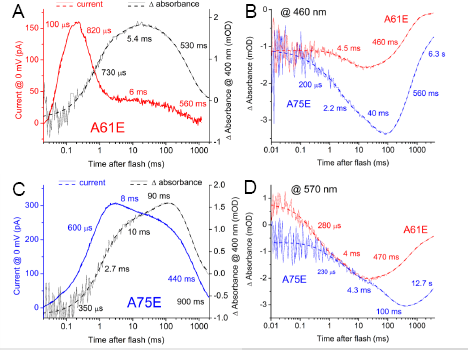
Temporal comparison of the laser flash-induced photocurrents and absorbance changes in the A61E and A75E mutants. (**A** and **C**) Photocurrent traces (red and blue, left axis) and absorbance changes (black, right axis) monitored at 400 nm recorded in response to laser flash excitation from the A61E mutant (A) and A75E mutant (B). (**B** and **D**) Flash-induced absorbance changes monitored in the A61E (red) and A75E (blue) mutants at 460 nm (B) and 570 nm (D).

The maxima of the absorption spectra of the purified A61E and A75E mutants are identical at 511 nm, and differ only 4 nm from the wild type (Figure 6 – figure supplement 1). Therefore neither glutamate residue introduced by mutation directly interacts with the chromophore site, as expected from their location > 9 Å from the Schiff base. Our interpretation is that the influence of these mutations on the photocycle is likely due to changes in the structure of the electric field in the photoactive site that influence the pK_a_s of the involved residues.

In summary, the results of our comparative analysis of laser-flash induced photocurrents and absorbance changes show that introducing a negative charge on the opposite sides of the photoactive site by mutation causes opposite changes in the channel kinetics and rectification, as is observed in the wild type upon shifting of the membrane potential in the opposite directions.

Comparative analysis of the photochemical reaction cycle transitions and photocurrent kinetics strongly indicates that the only open-channel conformer is the L intermediate, with possibly a small contribution from a red-shifted intermediate accompanying the L species (Sineshchekov et al. 2016). Depletion of L proceeds in two kinetics phases: a fast reversible L⇔M transition and a slower irreversible depletion of M and hence L. Application of an external electric field has opposite effects on fast and slow closing of the channel. The rate and contribution of the fast closing component increase and those of the slow closing component decrease (and vice versa, depending on the sign of the external field). We observed similar opposite effects upon electrostatic perturbation of the photoactive site by glutamate placed on the extracellular side versus the cytoplasmic side of the photoactive site. Our interpretation is that the fast and slow channel closing processes, kinetically correlated with the two phases of L dissipation, involve intramolecular charge displacements in opposite directions, which would explain opposite rectification of the two closing phases.

## Materials and Methods

### Expression of GtACR1 mutants in mammalian cells and patch clamp recording

For mammalian expression a DNA polynucleotide encoding the transmembrane domain (residues 1-295) of *Gt*ACR1 optimized for human codon usage was cloned into the pcDNA3.1 vector (Life Technologies, Grand Island, NY) in frame with an EYFP tag. Mutations were introduced using a QuikChange XL site-directed mutagenesis kit (Agilent Technologies, Santa Clara, CA) and verified by DNA sequencing. Characterization of *Gt*ACR1 mutants was performed using whole-cell photocurrent recording as previously described (Govorunova et al. 2015). HEK293 (human embryonic kidney) cells were transfected using the ScreenFectA transfection reagent (Waco Chemicals USA, Richmond, VA). All-*trans*-retinal (Sigma, St. Louis, MO) was added at the final concentration 4 µM immediately after transfection. Photocurrents were recorded 48-72 h after transfection in whole-cell voltage clamp mode at room temperature (25°C) with an Axopatch 200B amplifier (Molecular Devices, Union City, CA) and digitized with a Digidata 1440A using pClamp 10.7 software (both from Molecular Devices). Continuous light pulses were provided by a Polychrome V light source (T.I.L.L. Photonics GMBH, Grafelfing, Germany) at 15 nm half-bandwidth in combination with a mechanical shutter (Uniblitz Model LS6, Vincent Associates, Rochester, NY; half-opening time 0.5 ms). The maximal quantum density at the focal plane of the 40× objective measured with a piezo detector was 7.7 mW mm^-2^at 515 nm. Laser excitation was provided by a Minilite Nd:YAG laser (532 nm, pulsewidth 6 ns, energy 12 mJ; Continuum, San Jose, CA). A laser artifact measured with a blocked optical path was digitally subtracted from the recorded traces. For further analysis, the signals were logarithmically averaged with a custom-created computer algorithm. Patch pipettes with resistances of 2-5 MΩ were fabricated from borosilicate glass and filled with the following solution (in mM): KCl 126, MgCl_2_2, CaCl_2_0.5, EGTA 5, HEPES 25, and pH 7.4. The standard bath solution contained (in mM): NaCl 150, CaCl_2_1.8, MgCl_2_1, glucose 5, HEPES 10, pH 7.4. A 4 M KCl bridge was used in all measurements. Series resistance was periodically checked during recording, and measurements showing >20% increase were discarded. The holding potential values were corrected for liquid junction potentials calculated using the Clampex built-in LJP calculator (Barry 1994). Curve fitting and data analysis were performed using OriginPro 2016 software (OriginLab Corporation, Northampton, MA).

Batches of culture were randomly allocated for transfection with a specific mutant; no masking (blinding) was used. Individual transfected HEK293 cells were selected for patching by inspecting their tag fluorescence; non-fluorescent cells were excluded. Cells for which we could not establish a gigaohm seal or for which a gigaohm seal was lost during recording were excluded from measurements. Current traces recorded from the same cells upon repetitive light stimulation were considered as technical replicates; results obtained from different individual cells were considered as biological replicates. In experiments in experiments with continuous light pulses, a single trace was recorded in each cell; in experiments with laser excitation, 5-10 technical replicates were averaged to yield a single mean trace for each cell. The baseline measured before illumination was subtracted using Clampfit software (a subroutine of pClamp). The same software was used to measure the peak current amplitude with a cursor. The raw data obtained in individual cells are shown as open diamonds and listed in the corresponding source data tables. The sample size was estimated from previous experience and published work on a similar subject, as recommended by the NIH guidelines (Dell et al. 2002). No outliers were excluded from calculations. Normality of the data was not assumed, and therefore non-parametric statistical tests were used as implemented in OriginPro 2016 software; p values > 0.01 were considered not significant. The results of statistical hypothesis testing are given in Figure 2 source data file. When no specific statistical hypothesis was tested, descriptive statistical analysis was applied, and the results are presented as the mean values and SD or SE, as described in the figure legends.

### Expression of GtACR1 mutants in Pichia pastoris, *absorption spectroscopy and flash photolysis*

For expression in *Pichia*, the polynucleotides encoding *Gt*ACR1 mutants were fused in frame with a C-terminal eight-His tag and subcloned into the pPIC9K vector (Invitrogen) between EcoRI and NotI sites. The resultant plasmids were verified by DNA sequencing, linearized with SalI and used to transform P. pastoris strain SMD1168 (*his4, pep4*) by electroporation, as described earlier (Sineshchekov et al. 2016). The transformants were first screened for their ability to grow on histidine-deficient medium, and second, for their geneticin resistance. Up to eight single colonies that grew on 4 mg/ml geneticin were screened by small-scale cultivation, and clones of the brightest pink color were selected for further use.

For protein purification, a starter culture was inoculated into buffered complex glycerol medium until A600 reached 4–8, after which the cells were harvested by centrifugation at 5000 rpm and transferred to buffered complex methanol medium supplemented with 5 μM all-trans retinal (Sigma Aldrich). Expression was induced by the addition of 0.5% methanol. After 24-30 h, the cells were harvested and disrupted in a bead beater (BioSpec Products, Bartlesville, OK) in buffer A (20 mM sodium phosphate, pH 7.4, 100 mM NaCl, 1 mM EDTA, 5% glycerol). Cell debris and unbroken cells were removed by centrifugation at 5000 rpm. Membrane fragments were collected by ultracentrifugation at 40,000 rpm in a Ti45 rotor, resuspended in buffer B (20 mM Hepes, pH 7.4, 300 mM NaCl, 5% glycerol) and solubilized by incubation with 1.5% dodecyl maltoside (DDM) for 1.5 h or overnight at 4°C. Non-solubilized material was removed by ultracentrifugation at 50,000 rpm in a TLA-100 rotor. The supernatant was mixed with nickel-nitrilotriacetic acid agarose beads (Qiagen, Hilden, Germany) and loaded on a column. After washing the column thoroughly with buffer C (20 mM Hepes, pH 7.4, 300 mM NaCl, 5% glycerol, 0.02% DDM) containing 20 mM and 40 mM imidazole, the protein was eluted with buffer C containing 300 mM imidazole. The pigment was concentrated and imidazole was removed by repetitive washing with imidazole-free buffer C using YM-10 centrifugal filters (Amicon, Billerica, MA).

Absorption spectra of purified pigments in the UV-visible range were recorded on a Cary 4000 spectrophotometer (Varian, Palo Alto, CA). pH titration was carried out by the addition of small volumes of 1 M NaOH, 0.5 M Tris (pH 10), 0.5 M citric acid, or 1 M HCl. Light-induced absorption changes were measured with a laboratory-constructed crossbeam apparatus. Excitation flashes (532 nm, 6 ns, up to 40 mJ) were provided by a Surelite I Nd-YAG laser (Continuum, Santa Clara, CA). Measuring light was from a 250-W incandescent tungsten lamp combined with a McPherson monochromator (model 272, Acton, MA). A Hamamatsu Photonics (Bridgewater, NJ) photomultiplier tube (model R928) was protected from excitation laser flashes by a second monochromator of the same type and additionally with 12-nm bandwidth interference filters (Oriel Instruments, Stratford, CT). Signals were amplified by a low noise current amplifier (model SR445A; Stanford Research Systems, Sunnyvale, CA) and digitized with a GaGe Octopus digitizer board (model CS8327, DynamicSignals LLC, Lockport, IL), maximum sampling rate 50 MHz. The time interval between excitation flashes was 20 s, and up to 100 sweeps were averaged for each wavelength. Data analysis was performed with pClamp 10.7 (Molecular Devices, Union City, CA) and OriginPro 7 (OriginLab, Northampton, MA) software. Logarithmic filtration of the data was performed using the GageCon program provided by Dr. L.S. Brown (University of Guelph, Guelph, Canada).

### Other methods

Homology models of the *Gt*ACR1 mutants were created by I-TASSER suite (Yang et al. 2015) using the wild type *Gt*ACR1 structure (6EDQ) as the template. PyMOL software (http://www.pymol.org) was used for visualization and analysis of molecular structures. The intramolecular tunnel was identified by CAVER software (Kozlikova et al. 2014). Sequence logos were created using WebLogo algorithm (Crooks et al. 2004).

### Cell lines

Only a commercially available cell line authenticated by the vendors (HEK293 from ATCC) was used in this study; no cell lines from the list of commonly misidentified cell lines were used. The absence of mycoplasma contamination was verified by Visual-PCR mycoplasma detection kit (GM Biosciences, Frederick, MD).

## Acknowledgements

This work was supported by National Institutes of Health Grant R01GM027750, the Hermann Eye Fund, and Endowed Chair AU-0009 from the Robert A. Welch Foundation to J.L.S.

## Additional information Competing interests

E.G.G., O.A.S., J.L.S., and The University of Texas Health Science Center at Houston have filed patent applications that relate to ACRs (PCT application PCT/US2016/023095, titled “Compositions and Methods for Use of Anion Channel Rhodopsins”).

## Additional files

### Figure supplements and their legends

**Figure 1 – figure supplement 1.**
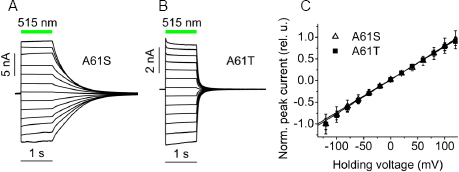
The current traces (A & B) and current-voltage relationships (C) for A61S and A61T mutants of *Gt*ACR1. (**A** and **B**). The current traces were recorded from HEK293 cells in response to 1-s light pulse (7.7 mW mm^-2^), the duration of which is shown as a green bar. The holding voltage was varied in 20-mV increments from -120 mV (the bottom trace) to 120 mV (the top trace). (**C**) The data points are mean values ± SE normalized at the value measured at -120 mV (n = 6 cells for each mutant). The lines are linear approximations.

**Figure 2 – figure supplement 1.**
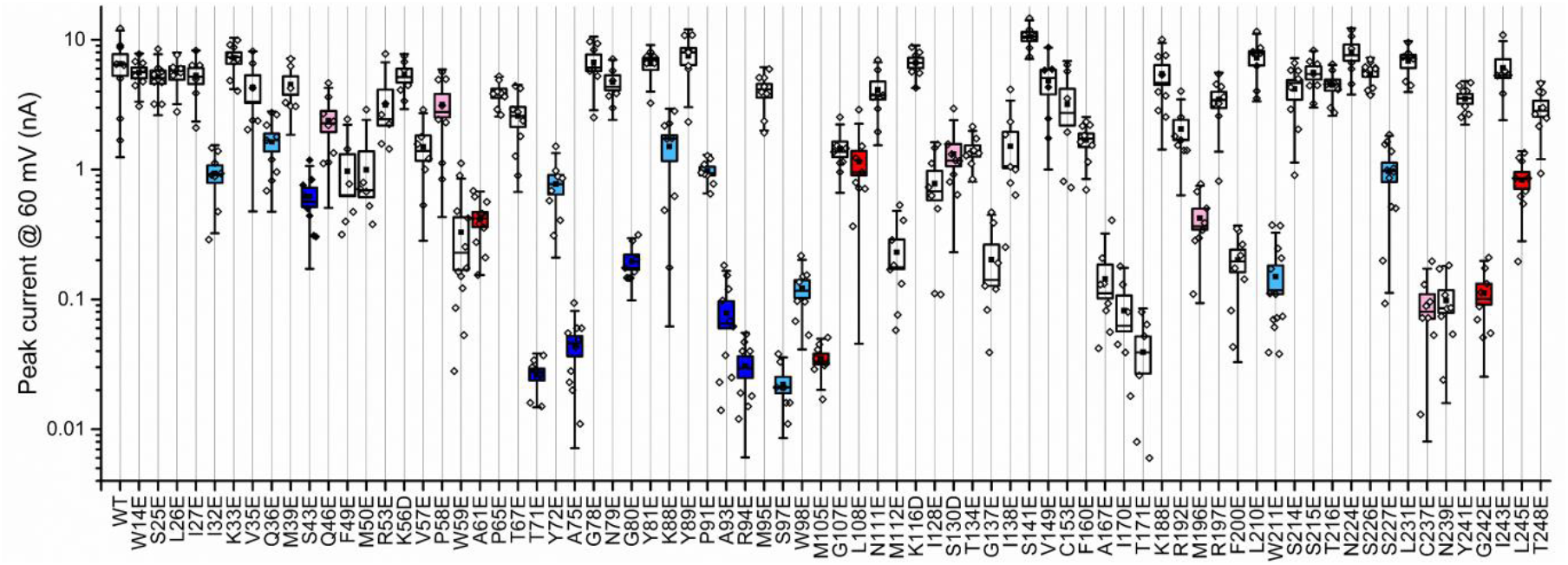
Peak current amplitude recorded at 60 mV holding voltage. Photocurrents were recorded in response to 1-s pulses of 515 nm light with 131 and 156 mM Cl-in the pipette and bath, respectively (for other solution components see Methods). The black squares, mean; line, median; box, SE; whiskers, SD; empty diamonds, raw data recorded from individual cells

**Figure 2 – figure supplement 2.**
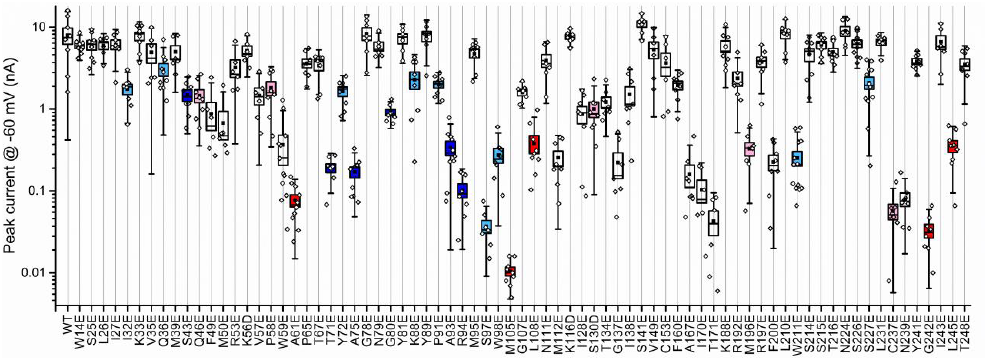
Peak current amplitude recorded at -60 mV holding voltage. Photocurrents were recorded in response to 1-s pulses of 515 nm light with 131 and 156 mM Cl-in the pipette and bath, respectively (for other solution components see Methods). The black squares, mean; line, median; box, SE; whiskers, SD; empty diamonds, raw data recorded from individual cells.

**Figure 2 – figure supplement 3.**
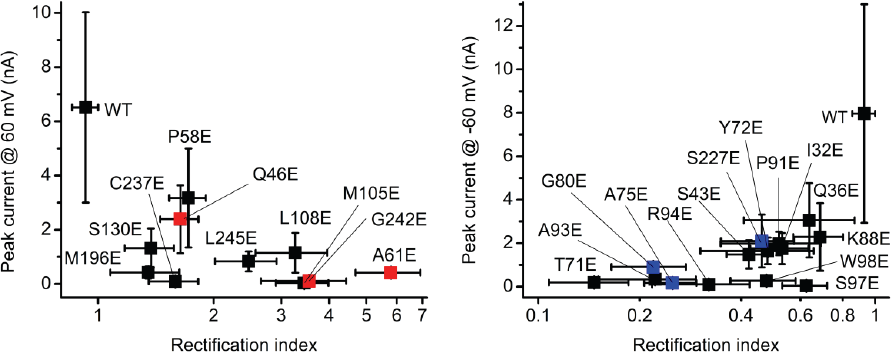
The dependence of the peak current amplitude at 60 and -60 mV on the rectification index measured in outwardly (A) and inwardly (B) rectifying *Gt*ACR1 mutants. The data points are the mean values ± SD (n = 6-10 cells; the exact number of cells tested for each variant is given in Figure 2 source data file).

**Figure 3 – figure supplement 1.**
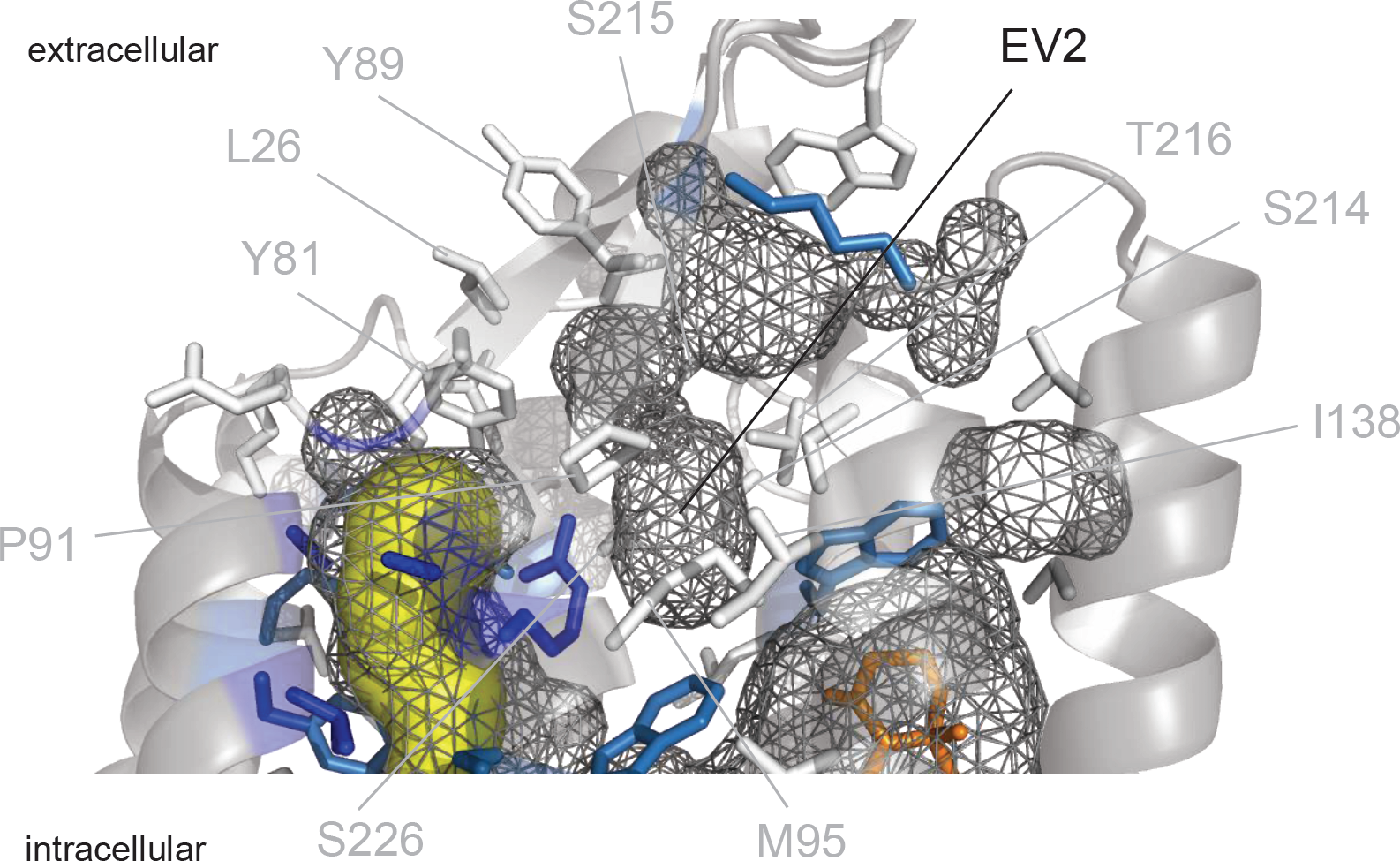
The second extracellular vestibule in *Gt*ACR1 (EV2). The side chains of contributing residues are color-coded according to the rectification index (RI) measured in their mutants, as defined in the Figure 3 legend. The intramolecular tunnel is shown in yellow, and the intramolecular surface, as grey mesh.

**Figure 4 – figure supplement 1.**
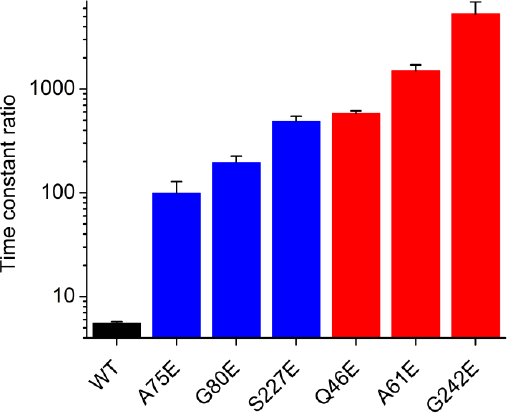
The ratio of the time constants (τ of the slow phase divided by τ of the fast phase) of the photocurrent decay recorded upon laser flash excitation. The time constants were derived by multiexponential fitting of the experimental current traces, as shown in Figure 4 in the main text. The data points are the mean values ± SE (n = 3-4 cells).

**Figure 4 – figure supplement 2.**
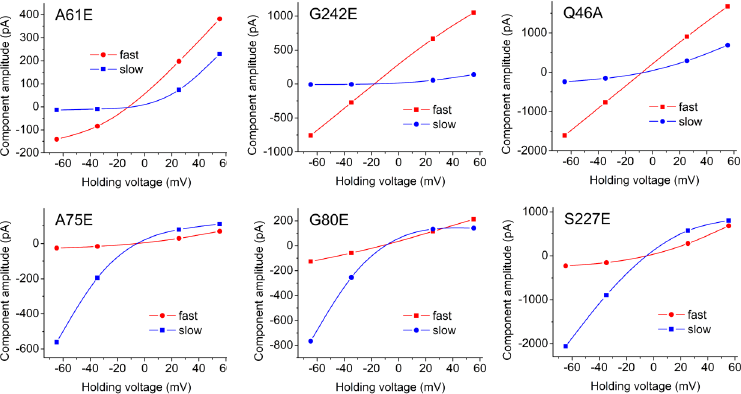
The dependence of the amplitude of the fast decay component on the holding voltage. The data points are the values obtained by multiexponential fitting of the current traces recorded upon laser flash excitation from representative cells for each mutant.

**Figure 5 – figure supplement 1.**
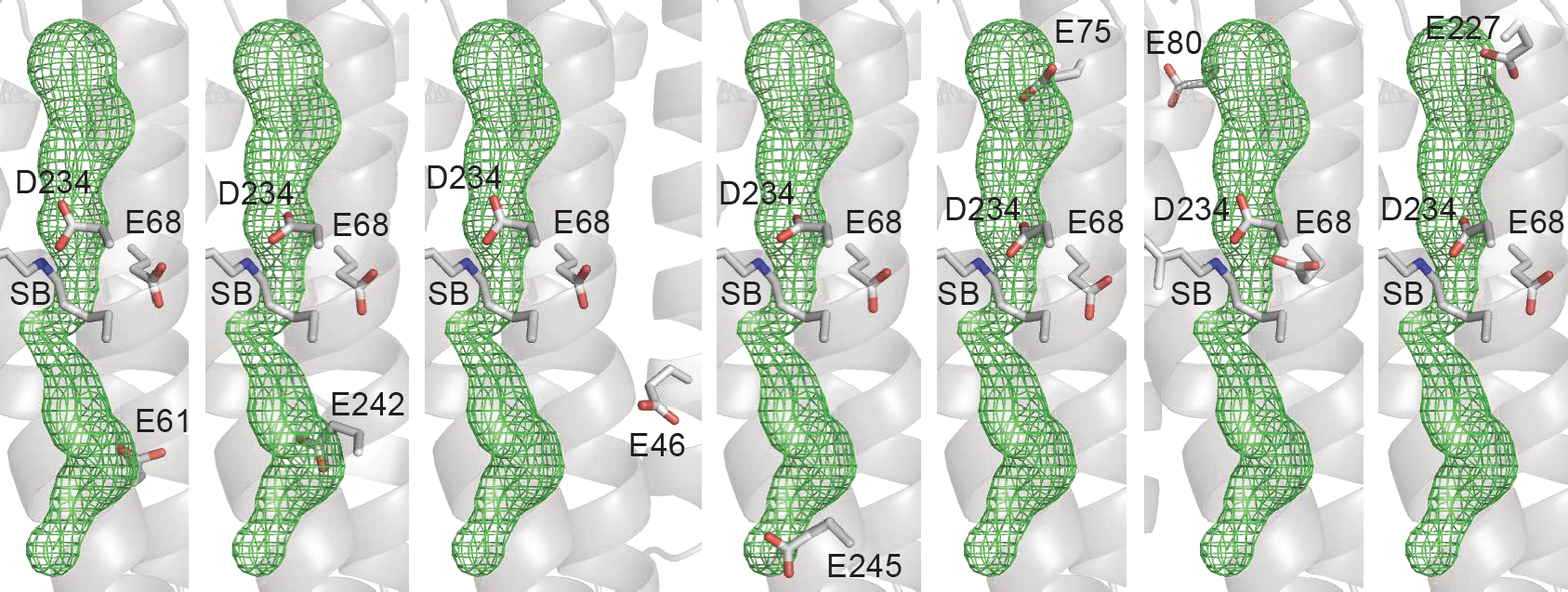
The homology models of the rectifying *Gt*ACR1 mutants. The models were created using the crystal structure of wild-type *Gt*ACR1 (6EDQ). The intramolecular tunnel is shown as green mesh, the side chains of the mutated residues and the photoactive center carboxylates are shown as sticks.

**Figure 5 – figure supplement 2.**
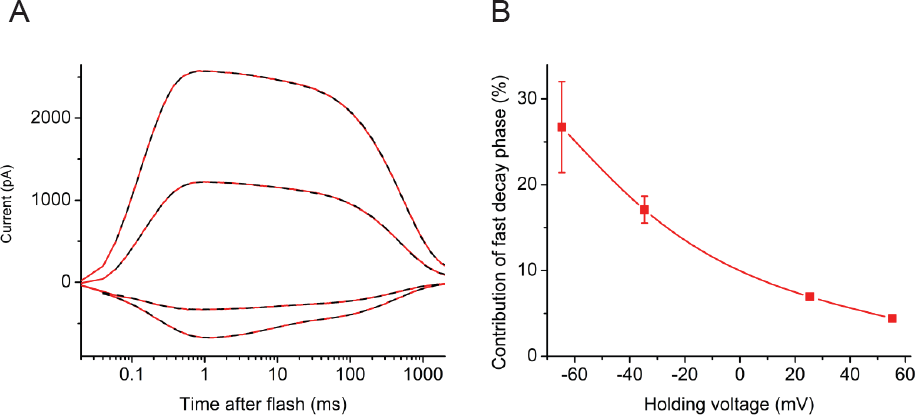
Kinetic analysis of photocurrents from the *Gt*ACR1_L245E mutant. (**A**) Photocurrent traces recorded in response to laser flash excitation at -60, -30, 30 and 60 mV at the amplifier output (bottom to top). The solid red lines show the experimental data, the dashed black lines, and their multiexponential fits. (**B**) The voltage dependence of the contribution of the fast decay phase.

**Figure 6 – figure supplement 1.**
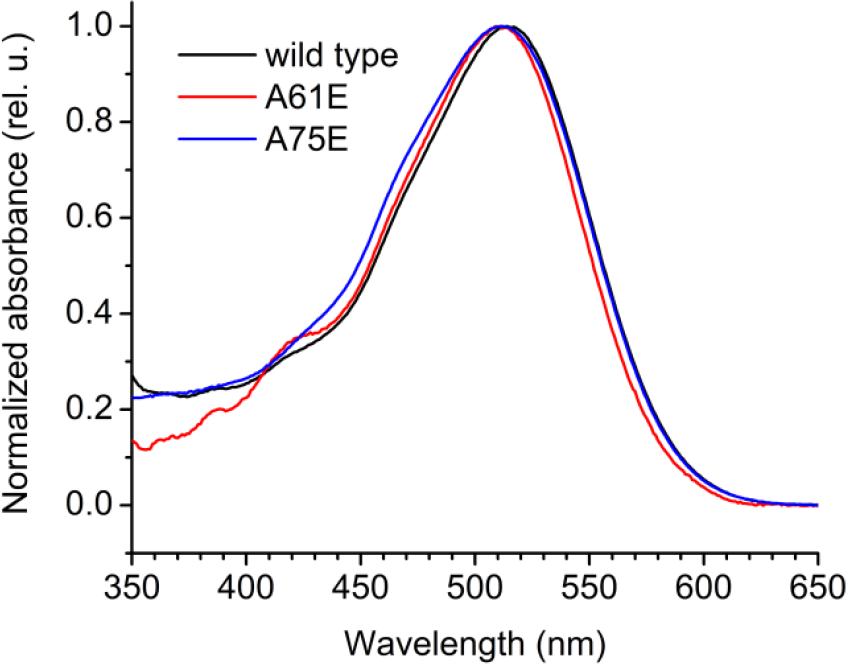
The absorption spectra of purified wild type *Gt*ACR1 and its mutants. The proteins were produced in *Pichia pastoris* and purified in non-denaturing detergent as described in Methods.

## References

Barry PH. 1994. Jpcalc, a software package for calculating liquid junction potential corrections in patch-clamp, intracellular, epithelial and bilayer measurements and for correcting junction potential measurements. J Neurosci Methods 51:107–116.

Crooks GE, Hon G, Chandonia JM, and Brenner SE. 2004. Weblogo: A sequence logo generator. Genome Res 14:1188–1190.

Dell RB, Holleran S, and Ramakrishnan R. 2002. Sample size determination. ILAR J 43:207–213.

Govorunova EG, Sineshchekov OA, Liu X, Janz R, and Spudich JL. 2015. Natural light-gated anion channels: A family of microbial rhodopsins for advanced optogenetics. Science 349:647–650.

Kato HE, Kim YS, Paggi JM, Evans KE, Allen WE, Richardson C, Inoue K, Ito S, Ramakrishnan C, Fenno LE, Yamashita K, Hilger D, Lee SY, Berndt A, Shen K, Kandori H, Dror RO, Kobilka BK, and Deisseroth K. 2018. Structural mechanisms of selectivity and gating in anion channelrhodopsins. Nature 561:349–354.

Kato HE, Zhang F, Yizhar O, Ramakrishnan C, Nishizawa T, Hirata K, Ito J, Aita Y, Tsukazaki T, Hayashi S, Hegemann P, Maturana AD, Ishitani R, Deisseroth K, and Nureki O. 2012. Crystal structure of the channelrhodopsin light-gated cation channel. Nature 482:369–374.

Kim YS, Kato HE, Yamashita K, Ito S, Inoue K, Ramakrishnan C, Fenno LE, Evans KE, Paggi JM, Dror RO, Kandori H, Kobilka BK, and Deisseroth K. 2018. Crystal structure of the natural anion-conducting channelrhodopsin *gt*acr1. Nature 561:343–348.

Kozlikova B, Sebestova E, Sustr V, Brezovsky J, Strnad O, Daniel L, Bednar D, Pavelka A, Manak M, Bezdeka M, Benes P, Kotry M, Gora A, Damborsky J, and Sochor J. 2014. Caver analyst 1.0: Graphic tool for interactive visualization and analysis of tunnels and channels in protein structures. Bioinformatics 30:2684–2685.

Li H, Huang CY, Govorunova EG, Schafer CT, Sineshchekov OA, Wang M, Zheng L, and Spudich JL. 2019. Crystal structure of a natural light-gated anion channelrhodopsin. Elife 8:e41741.

Mahn M, Gibor L, Patil P, Cohen-Kashi Malina K, Oring S, Printz Y, Levy R, Lampl I, and Yizhar O. 2018. High-efficiency optogenetic silencing with soma-targeted anion-conducting channelrhodopsins. Nat Commun 9:4125.

Messier JE, Chen H, Cai ZL, and Xue M. 2018. Targeting light-gated chloride channels to neuronal somatodendritic domain reduces their excitatory effect in the axon. eLife 7:e38506.

Mohammad F, Stewart JC, Ott S, Chlebikova K, Chua JY, Koh TW, Ho J, and Claridge-Chang A. 2017. Optogenetic inhibition of behavior with anion channelrhodopsins. Nat Methods 14:271–274.

Oda K, Vierock J, Oishi S, Rodriguez-Rozada S, Taniguchi R, Yamashita K, Wiegert JS, Nishizawa T, Hegemann P, and Nureki O. 2018. Crystal structure of the red light-activated channelrhodopsin chrimson. Nat Commun 9:3949.

Sineshchekov OA, Govorunova EG, Li H, and Spudich JL. 2015. Gating mechanisms of a natural anion channelrhodopsin. Proc Natl Acad Sci USA 112:14236–14241.

Sineshchekov OA, Li H, Govorunova EG, and Spudich JL. 2016. Photochemical reaction cycle transitions during anion channelrhodopsin gating. Proc Natl Acad Sci USA 113:E1993–2000.

Volkov O, Kovalev K, Polovinkin V, Borshchevskiy V, Bamann C, Astashkin R, Marin E, Popov A, Balandin T, Willbold D, Buldt G, Bamberg E, and Gordeliy V. 2017. Structural insights into ion conduction by channelrhodopsin 2. Science 358:eaan8862.

Wilson DE, Scholl B, and Fitzpatrick D. 2018. Differential tuning of excitation and inhibition shapes direction selectivity in ferret visual cortex. Nature 560:97–101.

Yang J, Yan R, Roy A, Xu D, Poisson J, and Zhang Y. 2015. The i-tasser suite: Protein structure and function prediction. Nat Methods 12:7–8.

